# The utility and caveat of split-GAL4s in the study of neurodegeneration

**DOI:** 10.1101/2022.12.06.519338

**Authors:** Luca Stickley, Emi Nagoshi

## Abstract

Parkinson’s disease (PD) is the second most common neurodegenerative disorder, afflicting over 1% of the population of age 60 years and above. The loss of dopaminergic (DA) neurons is the substantia nigra pars compacta (SNpc) is the primary cause of its characteristic motor symptoms. Studies using *Drosophila melanogaster* and other model systems have provided much insight into the pathogenesis of PD. However, little is known why certain cell types are selectively susceptible to degeneration in PD. Here we describe an approach to identify vulnerable subpopulations of neurons in the genetic background linked to PD in *Drosophila*, using the split-GAL4 divers that enable genetic manipulation of a small number of defined cell population. We identify a subtype of DA neurons selectively vulnerable in the model of the *leucine-rich repeat kinase 2* (*LRRK2*)-linked familial PD, demonstrating the utility of this approach. We also show an unexpected caveat of the split-GAL4 system in aging-related research: the age-dependent increase in the number of GAL4-labeled cells.

## Introduction

*Drosophila melanogaster* has emerged as a powerful model system to study neurodegenerative disorders, owing to the conservation of genetic programs and fundamental neurobiology between flies and humans, as well as its wealth of genetic tools [1]. A prime example is the research on Parkinson’s disease (PD), which is characterized by the loss of dopaminergic (DA) neurons in the substantia nigra pars compacta (SNpc) and the resulting locomotor impairments. DA neuron loss is frequently associated with the accumulation of intracytoplasmic inclusions mainly composed of the aggregates of *α*-synuclein protein, termed Lewy bodies (LBs), and those in the neurites, termed Lewy neurites (LNs) [2]. Discoveries of mutations linked to familial PD have led to the generation of numerous PD models in flies, which have made substantial contributions in understanding pathophysiology and pathogenic mechanisms of PD [3,4].

Understanding pathogenesis of PD requires the identification and characterization of vulnerable neuronal populations. While PD is increasingly recognized as a multisystem disorder, its pathology is observed nevertheless in restricted cell types. Postmortem analysis of PD patients tissues have shown the appearance of LBs and LNs in peripheral nervous system, including the enteric neurons and the autonomic nervous system, as well as in the multiple regions of the brainstem [5]. The LB and LN pathology is also observed in neocortex in the patients of advanced stages [6]. Notably, within the midbrain DA neuron population, the tegmental area (VTA) and the retrorubral area (RRA) are affected to a much lesser extent than the SNpc [5], suggesting that cell-type difference has a non-negligible role in the pathogenesis. It remains unclear whether specific subtypes of SNpc DA neurons are selectively vulnerable in PD, although the advent of single-cell RNA-sequencing technology is closing this knowledge gap [7].

Here we describe an approach to identify vulnerable subpopulations of DA neurons in the genetic background linked to PD in *Drosophila*. The method takes advantage of the split-GAL4 drivers, which enable the expression of transgenes in a small number of defined subsets of cells [8–10]. We show the utility of this powerful system by identifying a subtype of DA neurons selectively vulnerable in the model of the *leucine-rich repeat kinase 2* (*LRRK2*)-linked familial PD. We also show an unexpected and important caveat of this system for ageing-related research: the age-dependent increase in the number of cells labeled by the split-GAL4.

## Materials and Methods

### Drosophila culture and strains

Flies were raised on standard cornmeal-agar food at 25°C in a 12h: 12h light-dark cycle and under controlled humidity. Following fly strains were obtained from the Bloomington *Drosophila* stock center (BDSC): *20XUAS-6XGFP* (Bl #52261), MB316B (Bl #68317), MB032B (Bl #68302), MB056B (Bl #68276), MB109B (Bl #68261), MB025B (Bl #68299). *UAS-Lrrk^I1915T^* was described previously [11,12].

### Immunohistochemistry

Immunostaining was performed as previously described in [11]. Briefly, fly heads were fully dissected and fixed in 400mL of 4% paraformaldehyde (PFA), 0.3% Triton X-100 for 20 min on ice. Once fixation was completed, three quick washes followed by three 20-min washes were performed in PBS, 0.3% Triton X-100. Blocking was achieved with 1 h incubation in 5% normal goat serum (NGS), PBS, 0.3% Triton X-100 at room temperature on a rocking platform. Brains were incubated with the primary antibodies for two nights at 4°C on a rocking platform. Brains were then washed and incubated with the secondary antibodies overnight at 4°C. Vectashield (Vector Laboratories) was used as slide mounting medium. Antibodies used in this study and dilutions were as follows. Mouse anti-nc82 (mouse, Developmental Studies Hybridoma Bank, 1:100), Rabbit anti-GFP (G10362, Invitrogen. 1:500), Goat anti-Rabbit IgG Alexa 488 1:200 (A11008, Thermofisher, 1:200), and Goat anti-mouse IgG-Alexa 633 1:200 (mouse, A21052, Thermofisher, 1:200).

### Imaging and image analysis

Fly brains were scanned using Leica TCS SP5 confocal microscope, at 40x with 1.4x zoom Quantification of cell numbers was performed as described (Bou Dib et al. (2014); Tas et al. (2018)), using the cell counter plugin of Fiji (Fiji is just ImageJ) [13].

### Statistics and graphics

Statistics were performed in python with the help of several statistical packages. Comparison between two conditions was performed with the ttest_ind function of the python SciPy Stats package [14]. Two-tailed Student’s t-test for equal variance, and Welch’s t-test for unequal variance. Groups with more than one factor were compared with ANOVA with a Tukey’s HSD post-hoc test using the pairwise_tukeyhsd function of the python statsmodel package [15]. Statistical significance for all comparisons was set at p<0.05. Graphical representation of data distributions was performed according to the guidelines of raincloud plots with the PtitPrince python package [16].

## Results

We and others have previously shown the loss of dopaminergic neurons (DA) within the protocerebral anterior medial (PAM) cluster by genetic or pharmacological insults in adult *Drosophila* brains [11,17–19]. Although the number of PAM neuron loss observed so far varies between 15% and 80%, other DA clusters remain unaffected. These results are reminiscent of regionally restricted dopaminergic neurodegeneration described in early human PD cases, where substantia nigra pars compacta (SNpc) DA neurons are more susceptible to PD neurodegeneration than ventral tegmental area (VTA) DA neurons [20]. The underlying cause of this distinction between nuclei is unclear but much of the discussion revolves around morphological as well as electrophysiological differences [20].

To investigate potential morphological and electrophysiological differences between degenerating and non-degenerating DA neurons, we wanted to find and describe a susceptible PAM subpopulation. PAM neurons comprise approximately 20 subpopulations, projecting to different subdomains of the mushroom body (MB) [21]. Taking advantage of the split-GAL4 divers targeting different subpopulations, we sought to identify PAM neuron subtypes that are vulnerable in PD-linked genetic predispositions. To this end, we focused on a genetic model of *leucine-rich repeat kinase 2* (*LRRK2*)-linked familial PD, based on a targeted expression of a mutant form of *Drosophila Lrrk* (*Lrrk^11915T^*) [12]. The split-GAL4 lines function by using two promoter sequences, one regulating the expression of the DNA binding domain (DBD.GAL4), while the second the activator domain (AD.GAL4), which then combine to form a fully functional GAL4 [8]. We selected four lines targeting the PAM-β‘2 subdomain and one line targeting PAM-β‘1, because PAM neurons projecting to the MB β‘ lobe is required for startle-induced locomotion [22] and their functional impairments accounts for locomotor deficits in some PD models [23] (Table 1). Evaluation of the susceptibility was performed by expressing either *UAS-GFP* or *UAS-GFP* in combination *UAS-Lrrk^11915T^* with each respective driver, followed by dissection at the age day 35 and counting of GFP-positive neurons. Expression of *Lrrk^11915T^* with MB032B and MB109B caused no significant reduction in GFP-positive cell counts, while a significant decrease was observed with MB056B (Fig. 1A, D and E). No reduction in the MB025B-expressing cells but instead a significant increase was observed in the group expressing *Lrrk^11915T^* (Fig. 1A). These results suggest that MB056B positive but MB032B and MB109B negative cells, i.e. PAM-β‘2p neurons, are susceptible to LRRK2-induced neuronal loss.

**Figure 1.**
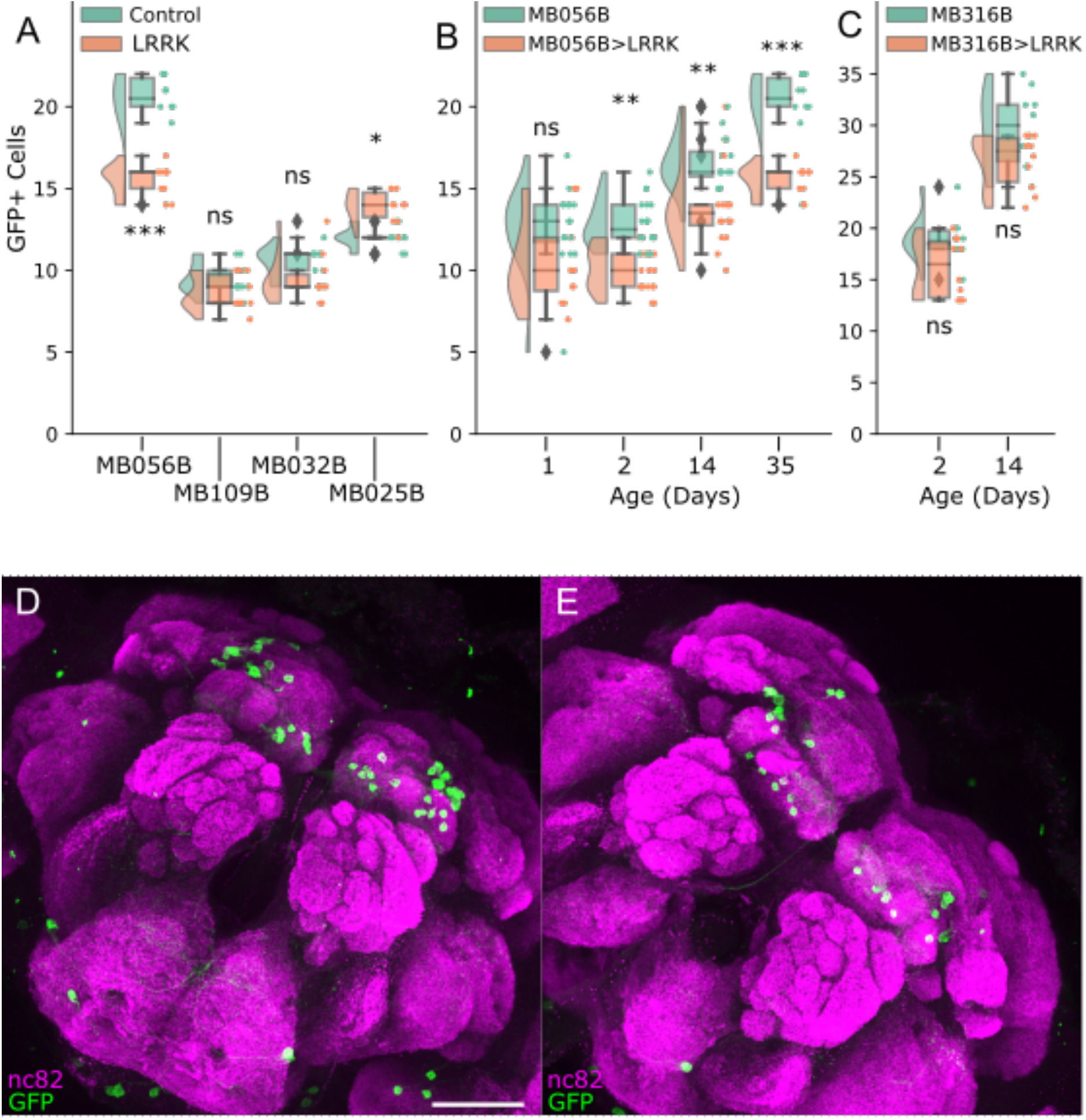
PAM-β’2 neurons show increased susceptibility to degeneration by *Lrrk^I1915T^*expression. (**A**). The number of GFP-positive cells in the flies expressing *GFP* with (LRRK) or without (Control) *Lrrk^11915T^* driven by the indicated split-GAL4 drivers. Day 35. Tukey HSD, MB056B p=0.001 (***) (n=10); MB109B p=0.6941 (n=10); MB032B p=0.5872 (n=10); MB025B p=0.0319 (*) (n=10). (**B**) The number of GFP-positive cells in MB056B control or MB056B>*Lrrk^11915T^* at days 1, 2, 14, and 35. The graph shows progressive increase in the both the number of neurons and disparity between treated and untreated groups. Statistics comparing treated and untreated for each time-point. Tukey HSD; Day 1 p=0.0864 (n=12-16), Day 2 p=0.0011 (**) (n=20), Day 14 p=0.0078 (**) (n=16), Day 35 p=0.001 (***) (n=10). (**C**) Number of GFP+ cell in MB316B control or MB056B>*Lrrk^I1915T^* at day 2 and 14. An increment in number of labelled neurons is also present in this split-GAL4. No comparison between control and *Lrrk^11915T^* groups is significant. Tukey HSD; Day 2 p=0.181 (n=10), Day 14 p=0.0843 (n=10). (**D** and **E**) MB056B (**D**) and MB056B>*Lrrk^11915T^* (**E**) at day 35. max projections showing neuropil staining by anti-NC82 and MB056B-labeled cells with anti-GFP. Scale bars = 50 μm.

**Table 1.** List of Split-GAL4 lines used and the PAM neuron subtypes targeted. (AD.Gal4) Activator domain inserted at site attP40 (2L) landing site, while DNA Binding Domain (DBD.Gal4) uses the attP2 (3L) landing site. Expression, PAM neuron subtypes targeted by each line, as described in [21].

| Name | AD.Gal4 (attP40) | DBD.Gal4 (attP2) | Expression |
| --- | --- | --- | --- |
| MB316B | R58E02 | R93G08 | PAM-$\beta'$2m, PAM-$\beta'$2p, PAM-γ4, PAM-γ4<γ1γ2 |
| MB032B | R30G08 | ple | PAM-$\beta'$2m |
| MB056B | R76F05 | R80G12 | PAM-$\beta'$2m, PAM-$\beta'$2p |
| MB109B | R76F05 | R23C12 | PAM-$\beta'$2a |
| MB025B | R24E12 | R52H01 | PAM-$\beta'$1ap, PAM-$\beta'$1m |

To continue investigating vulnerability of PAM subpopulations, it was necessary to determine if loss of dopaminergic neurons was solely developmental or increased with age. MB056B-labeled cells observed at day 1 and 2 showed promising results, with clearer separation between the control and *Lrrk^11915T^*groups in older flies (Fig. B). This trend continued at timepoints day 14 and day 35, but of grave concern was the ever-increasing number of GFP positive neurons in both control and *Lrrk^11915T^* groups. Although loss of neurons in MB316B-labeled population was not significant, the increase of GFP labeled cells was still present (Fig. C). Unfortunately, the issue of increasing GFP^+^ cell numbers by age in split-GAL4 lines is incompatible with the study of age-dependent loss of DA populations.

## Discussion

### Split-GAL4 incremental labeling

The number of cells expressing GFP driven by the MB056B and MB316B split-GAL4 drivers increases by age and does not plateau even between days 7 and 35 of age. Whereas classical GAL4s are expressed monolithically, split-GAL4s require the activator domain (AD.Gal4) and the DNA-binding domain (DBD.Gal4) to combine and perform its function. This is achieved by placing a leucine zipper (Zip) on a flexible linker at the N-terminus of Gal4-DBD.Gal4 (Zip-) and the C-terminus of p65-AD.Gal4 (Zip+) [8,9]. Although age-dependent increase in the activity of one or both of the promoters expressing the split-GAL4 components could result in the incremental GAL4 reconstitution, it seems unlikely that activity of involved promoters does not reach a plateau until day 35, if not impossible. As MB056B and MB316B do not share either AD.Gal4 or DBD.Gal4 promoters, such a mechanism would be independent of the inserted sequences, for example via changes in chromatin accessibility by age. Instead, low turnover of reconstituted GAL4 or GAL4-promoter complex may be a key contributor of the incremental labeling. The binding of monolithic GAL4 to the promoter has a short half-life *in vivo* [24]. It is worth investigating if reconstitution of GAL4 by heterodimeric leucine zippers increases GAL4 stability or changes turnover rate of GAL4-promoter binding. This feature, if indeed the case, would be an advantage in some application.

This study examined only two split-GAL4 lines for age-dependent expression changes, and it remains to be seen whether the finding is generalized. Yet, our results call for caution in using the split-GAL4 system. When using split-GAL4s, the comparison between groups at the same age will be legitimate, whereas comparisons between different age groups would not produce meaningful results.

### Functional and morphological vulnerability

Although the artifact of split-GAL4 system did not allow continued investigations of this topic, the finding of possible functional susceptibility is still of note. We found that PAM DA neurons projecting to a specific region of the MB, β’2p subdomain, were more susceptible than others. 90% of DA neurons that project to the MB receive feedback of the MB output neurons (MBON) in the convergent zones (crepine, superior medial protocerebrum, superior intermediate protocerebrum, and superior lateral protocerebrum) [1], which in turn receive dendrites from the central complex, a region crucial for motor actions [1]. Scaplen et al. [26] has recently shown that the MBON-β‘2mp and MBON-γ5β‘2a have the highest number of projections to PAM neurons, while MBON-β‘1 had one of the lowest. Although it is unclear if the PAM neurons synapsed by these MBONs are the same that innervate them, the differences in the PAM-MB-MBON-PAM feedback loop could explain the increased loss of neurons in PAM-β’2 neurons.

Work by Zhao et al. [27] has shown that *LRRK2* differs in its interactome depending on tissue, with a clear shared features found when in the putamen, caudate and nucleus accumbens. This finding encourages the studies of PAM-β‘2 susceptibility even with the hindrance of incremental labeling by split-GAL4s. Future work on PAM-β‘2 neurons might benefit from quantifying key morphological parameters of mitochondria, such as mitochondrial density, biomass, integrity, motility, and localization.

## Acknowledgements

We thank the Bloomington *Drosophila* Stock Center for fly stocks. We thank Sean Sweeney for comments on the manuscript.

## Author contributions

LS and EN conceptualized the work. LS performed experiments and analyzed the data. LS and EN wrote the manuscript.

## Data availability statement

All the data are presented in the paper. The original image data are available upon request.

